# NGSTroubleFinder: A tool for detection and quantification of contamination and kinship across human NGS data

**DOI:** 10.1101/2025.01.31.635690

**Authors:** Samuel Valentini, Tecla Venturelli, Xavier Gallego, Laura Perez-Cano, Emre Guney

**Affiliations:** STALICLA Discovery and Data Science Unit, World Trade Center, Moll de Barcelona, Edif Este, 08039 Barcelona, Spain

## Abstract

**Summary:** Quality control is a fundamental but often neglected step in any NGS pipeline. Detecting issues like cross-sample contamination and sample swaps is essential to control the data integrity. Here, we present NGSTroubleFinder, a novel python tool to detect cross-sample contamination in human Whole-Genome and Whole-Transcriptome Sequencing data, sample swaps and mismatches between the reported and the inferred genetic and transcriptomic sexes. NGSTroubleFinder is implemented in Python and incorporates a custom-built parallelized pileup engine written in C. The tool reports extensive information on the samples both in textual and HTML format including key plots for easy interpretation of the results.

**Availability and Implementation:** NGSTroubleFinder is written in Python and C, and it can be easily installed with pip. The tool source code and the models are freely available on github (https://github.com/STALICLA-RnD/NGSTroubleFinder) and a containerized version is available on dockerhub (https://hub.docker.com/r/staliclarnd/ngstroublefinder).

## Introduction

Next generation sequencing (NGS) has revolutionized genomics, transcriptomics, and epigenomics by generating vast amounts of data that provide unprecedented insights into biological systems. However, the validity of these insights heavily depends on the integrity of the sequencing data, making quality control a fundamental step in NGS data analysis. Contamination in NGS samples, whether from exogenous sources or cross-sample contamination, is not uncommon due to the laborious experimental steps prone to human error. Such contamination can significantly skew results and lead to erroneous conclusions.

Tools like multiQC (Ewels *et al*. 2016) allow for fast and reliable identification of issues related to the technical quality of a sequencing experiment like read quality, duplication, and alignment statistics. A higher-than-expected ratio of unmapped reads, followed by alignment of some of these reads, is often sufficient to pinpoint potential viral and bacterial contamination (Nieuwenhuis *et al*. 2020; Selitsky *et al*. 2020; Chrisman *et al*. 2022). However, post-alignment quality control to identify issues like sample swaps and cross-sample contamination are often overlooked and not typically included in standard quality control procedures (Zhou *et al*. 2018).

Several tools analyzing allelic ratios, k-mer frequencies, or gene markers such as VerifyBamID2 (Zhang *et al*. 2020) and read_haps (Eggertsson and Halldorsson 2021) have been developed to identify and quantify contamination. Other tools like ContEst, Conpair, Vanquish, ART-DeCo, CleanCall, VICES, MICon, and CHARR have been benchmarked in (Chen *et al*. 2024). However, several of the aforementioned tools exhibited execution limitations, and none of them demonstrated applicability to RNA-sequencing data analysis.

On the other hand, tools like Somalier (Pedersen *et al*. 2020) and NGSCheckmate (Lee *et al*. 2017) are often employed to identify kinship and sample swaps. However, the integration of these tools into quality control pipelines is hindered by the need for installation of various tools with different requirements and the necessity for substantial technical expertise to effectively utilize and interpret the results.

Here, we present NGSTroubleFinder, a novel easy to use command line tool to detect and quantify sample cross-contamination and sample swaps in both Whole-Genome DNA Sequencing (WGS) and Whole-Transcriptomic RNA Sequencing (WTS) data from human samples (**Figure 1**). It aims to ensure the reliability of NGS data and the validity of subsequent biological and clinical interpretations. The command line tool can be seamlessly integrated in NGS data analysis pipelines, including but not limited to workflows written in Nextflow, and produces a detailed report that can be visualized interactively in any modern web browser. NGSTroubleFinder relies on a simple yet powerful linear regression model that leverages information on common variants to quantify likely cross contamination as well to offer insights on the swaps and kinship between samples. In addition to sample contamination and kinship estimation, NGSTroubleFinder can identify sex mismatches in which the biological sex inferred from the NGS data using variant and transcriptomic information is inconsistent with the assigned sex in the metadata.

**Figure 1.**
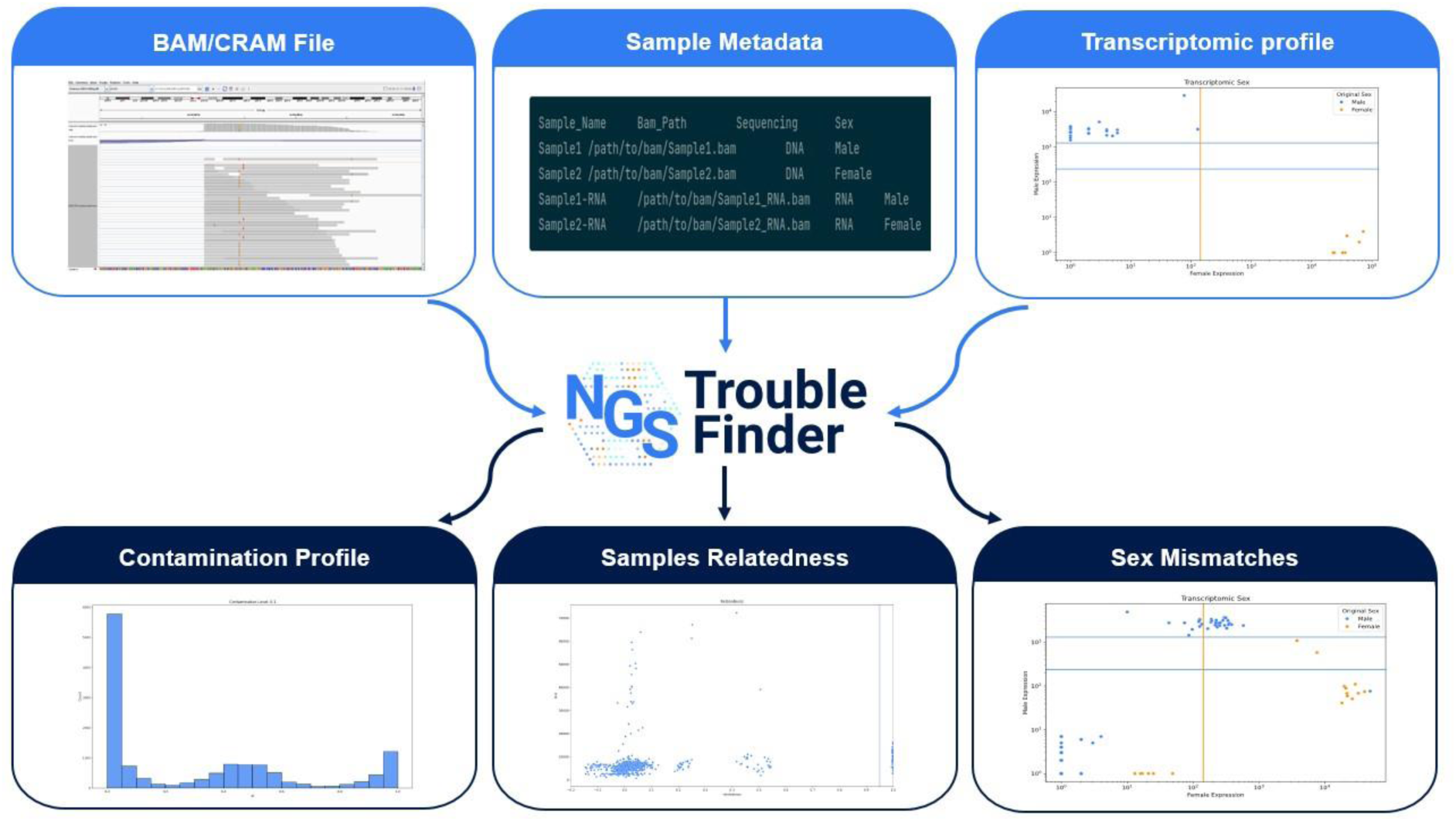
NGSTroubleFinder input and output. NGSTroubleFinder takes as input the BAM/CRAM files for a choort, the metadata of a choort and the transcriptomic profile of the sample (if available). It reports the contamination profile of the samples, their relatedness and if there are any mismatches on the reported sexes.

## Methods and Implementation

### Curated common variants

NGSTroubleFinder is based on a curated set of very common (MAF between 0.1 and 0.9 in the general population) human variants. Biallelic variants have been extracted from dbSNP (build id 151) (Sherry 2001) and filtered by keeping exonic variants, that are located at least 15 bases from a known INDEL and that are not located in hypervariable regions containing tandem repeats. The curated set of variants is included in the tool and it’s available in the github repository.

### Custom pileup engine, genotyping and haplotype detection

NGSTroubleFinder leverages a custom pileup engine written in C and based on the htslib (Bonfield et al., 2021) to compute a pileup of the curated set of variants. The pileup uses a strict quality approach considering a read only if the base quality is at least 30 and its mapping quality is greater than 1. Variants are genotyped using a heuristic approach based on the allelic fraction (AF), where REF and ALT correspond to reference and alternative alleles, respectively.

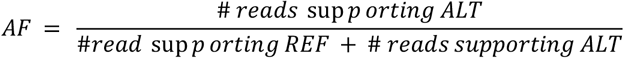

If the AF is less than 0.02 or greater than 0.98 the variant is genotyped as reference homozygous or alternative homozygous respectively. If the variant has an AF from 0.2 to 0.8 it is considered heterozygous. If the variant’s AF is outside those ranges, it is considered affected by noise and not called. Variant genotype is only utilized in kinship detection and not in contamination detection. The tool considers only variants with a coverage of at least 20 in all subsequent computation.

Secondly, close (i.e., less than 150 bases afar) high-quality pairs of variants are used to detect anomalies in the haplotypes following a methodology similar as read_haps (Eggertsson and Halldorsson 2021). In detail, if a read spans two known variants, only two of the four possible read combinations in a diploid individual (Reference-Reference, Reference-Alternative, Alternative-Reference, and Alternative-Alternative) should be observed. The region is flagged as an anomaly if three or more combinations are observed. A combination is considered observed if at least three non-duplicated reads are supporting the combination.

### Contamination estimation

Two linear regression models, one for WGS and one for WTS, have been trained on 50 samples (Table S1) randomly selected from the individual of European (EUR) ancestry in the 1000 Genome Project (1000GP) (Fairley *et al*. 2020). Those 50 samples have been considered contamination-free. From those 50 samples, contaminated samples were derived by mixing a controlled number of reads from another sample selected at random. Other 50 samples have been generated with the following contamination levels: 0.5%, 1%, 1.5%, 2%, 2.5%, 3%, 4%, 5%, 10%.

The test dataset has been generated using independent samples available internally at our institution. The WGS test dataset is based on 16 samples from 2 different batches which have been used to derive 15 samples per contamination level. The WTS dataset is based on 24 samples from a single batch which have also been used to derive 15 samples per contamination level.

The linear regression model uses three features:

- Number of SNPs with an allelic fraction between [0.05-0.15] or [0.85-0.95] normalized over the total number of SNPs (SNPs in tails)
- Number of SNPs with an allelic fraction between [0-0.02] or [0.98-1] normalized over the total number of SNPs (Homozygous SNPs)
- Number of regions supporting more than 2 haplotypes over the total number of available positions (Multiple Haplotypes)

The models are made available in Open Neural Network Exchange (ONNX) format in the github repository to facilitate findability, accessibility, interoperability, and reusability of these models in an easy, free and platform independent manner.

### Kinship identification

Kinship is identified by using high quality genotyped SNPs i,e., those that have a coverage depth of at least 20. Kinship estimation follows the model introduced in (Manichaikul *et al*. 2010), similarly to the tool Somalier (Pedersen *et al*. 2020).

The following metrics are introduced.

- ibs0 -- the number of sites where one sample is homozygous reference, and another is homozygous alternative
- ibs2 -- the number of sites where the samples have the same genotype
- sharedHets -- the number of sites where both samples are heterozygotes
- Het_i_ is the number of heterozygous site for sample i

Given two sample i,j their kinship is defined as:

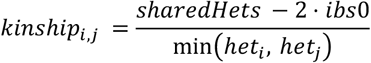

Two samples are considered replicates (or twins), if their kinship index is greater than 0.95. Two samples are considered related if the kinship goes from 0.15 to 0.95. Finally, they are considered unrelated for values less than 0.15.

### Sex mismatch identification

The sex of the sample is identified using two methodologies: variant based, and transcriptome based. In the variant based approach, variants on the X and Y chromosomes are used and the heterozygosity ratio (Samuels *et al*. 2016) of the X chromosome and total coverage of the variants on the Y chromosome are computed. A sample is considered female if it’s heterozygosity ratio on X is greater than 1 and the Y coverage is less than 10. Similarly, a sample is considered male if the heterozygosity is less than 1 and the Y coverage is greater than 10. Samples outside these ranges are flagged as anomalies. The transcriptome-based approach is based on the biomarkers reported in (Staedtler *et al*. 2013) using the integer read counts of the 4 male genes (RPS4Y1, EIF1AY, DDX3Y, KDM5D) and the female gene (XIST) whenever available (referred as discriminative genes hereafter). The expression level of these discriminative genes across individuals in the whole blood GTEx dataset (V8) (Lonsdale *et al*. 2013) is used to generate two reference distributions of read counts, one for females and one for males. For a given sample, first, the expression values of the discriminative genes on the Y chromosome are summed. Then, a sample is identified as female if the expression of XIST is greater than the 1^st^ percentile of the female reference distribution while the sum of the expression of discriminative genes on Y chromosome is less than the 99^th^ percentile of the female reference distribution. Conversely, if the expression level of discriminative genes on Y chromosome is greater than the 1^st^ percentile and the XIST expression is lower than the 99^th^ percentile in the male population, the sample is identified as male. If a sample is outside those ranges, it’s marked as an anomaly. If the expression levels for a sample are not available, the sample is marked as unknown.

### Implementation details

NGSTroubleFinder is a command-line tool implemented in Python and uses an internal, custom built pileup engine written in C (tested with Python 3.10.10, GCC 11.4.0 and libhts-dev version 1.13+ds-2build1) (Bonfield *et al*. 2021). It takes as input the path pointing to the output folder location and the path pointing to the metadata file containing the information on *(i)* sample name *(ii)* path pointing to the aligned sequencing file (i.e., BAM or CRAM), *(iii)* the type of sequencing (i.e., DNA or RNA), *(iv)* the expected sex for each sample. Optionally, a transcriptomic counts file containing integer read counts of each sample needs to be provided to enable biological sex inference from RNA. If the CRAM format is used for aligned sequencing files, the reference genome id also needs to be provided as input. The source code is freely available on github and a containerized version in docker is available on dockerhub.

### Quality control (QC) report

Extensive reports are produced about the inferred sex (Figure S1) kinship (Figure S2), and pileups (Figure S3). Additionally, per sample pileups are provided using the format used by PaCBAM (Valentini *et al*. 2019) and all the positions where haplotypes can be inferred are reported with the regions where multiple haplotypes are detected. Finally, a complete HTML interactive report is generated that can be used to explore the results.

### Benchmarking

We compared our results against the state-of-the-art tool VerifyBamID2 (Manichaikul *et al*. 2010) on our test dataset. We used the 1000GP 100k dataset to identify contamination in the WGS and the 1000GP 10k dataset for the WTS. Since both tools return a level of contamination,x their results can be compared against a ground truth using the mean absolute error metric. Overall, NGSTroubleFinder and VerifyBamID2 results are comparable, indicating a good generalization of our model (**Table 1**).

**Table 1.** Mean abs error of estimated contamination using NGSTroubleFinder and VerifyBamID2.

| Dataset | NGSTroubleFinder<br>Train dataset (abs error) | NGSTroubleFinder Test<br>dataset (abs error) | VerifyBamID2 Test<br>Dataset (abs error) |
| --- | --- | --- | --- |
| RNA-seq | 0.0084 | 0.0079 | 0.0083 |
| WGS | 0.0033 | 0.0066 | 0.0070 |

The absolute error among the different contamination classes shows that for the RNA Test dataset the results between NGSTroubleFinder and VerifyBamID2 are comparable. Interestingly, the average error is increasing with increasing contamination indicating that high levels of contamination are more difficult to estimate correctly by both tools (Table S2). The results for the DNA dataset are similar but VerifyBamID2 shows slightly lower error for low contamination levels than NGSTroubleFinder (Table S3).

## Discussion and limitations

We present NGSTroubleFinder a novel easy to use command line tool written in Python for NGS quality control. NGSTroubleFinder detects possible sample contamination in WGS and WTS, identifies possible sample swaps if at least two replicates of the same sample are available and kinship. It also infers the sex of the samples through a genomic and a transcriptomic method. It’s easy to use and it implements its pileup engine for fast computation.

The models the tool uses pose certain limitations, as they have been trained only on germline samples of European genetic background. Furthermore, the model performance can be limited if applied to cancer samples due to the presence of somatic mutations and copy number alterations and on WES due to the difference in coverage depth, but custom models for cancer/WES can be built and benchmarked. The model is also not suitable to detect inter-species contamination since it’s based on human variants. Also, we expect the contamination to be overestimated if the two admixed samples belonged to individuals from different ethnic groups. In the future, data that is being produced within the scope of recent initiatives such as “All of Us”(Bick *et al*. 2024) and “In Our DNA SC” (Allen *et al*. 2024) can be used to integrate population-specific variant information into the models.

Despite limitations inherent to lack of diversity in current human genetic data, NGSTroubleFinder addresses an important need towards assessing the reliability of NGS data from human samples that has widespread use in generating biological and clinical insights. The command line tool can be seamlessly integrated in NGS data analysis workflows and provides crucial information on the cross-sample contamination and swaps as well as familial relationships and sex mismatches between samples in a user-friendly report that can be visualized in a web browser in a platform independent manner.

Overall, we showed that the performance of NGSTroubleFinder in identifying sample contamination is comparable with state-of-the-art tools while providing extensive information on quality control including level of estimated contamination, identified sample swaps and family relationships and sex predicted based on NGS data. The tool is easy to use and is available on github and containerized on dockerhub.

## Acknowledgements

The authors wish to thank José Manuel Hidalgo López, Sergio Morales, Lynn Durham for valuable feedback and discussions, Anezka Zajicova for supporting generation of visuals in the manuscript and Hector Naranjo for computational infrastructure support.

