## Supplementary Information for "NGSTroubleFinder: A tool for detection and quantification of contamination and kinship across human NGS data"

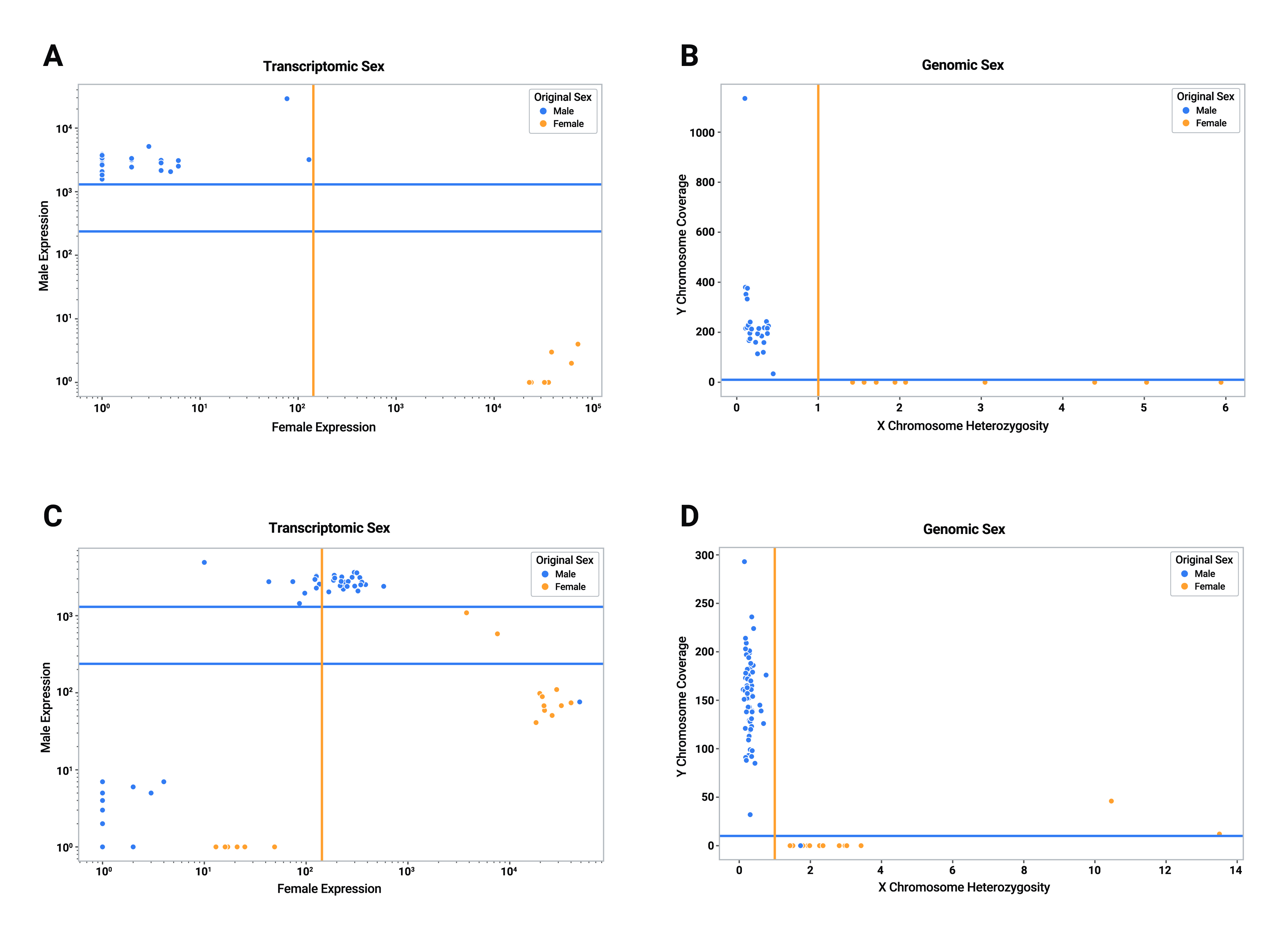

**Figure S1: Scatter plots generated using the tool for (A) transcriptomic sex identification and (B) genomic sex identification on the same dataset. Samples are colored by the provided sex while the lines are providing a guideline on how the samples are classified. (C) The scatter plot for transcriptomic identification generated using a contaminated dataset. Some samples are outside the classification thresholds and are marked as anomalies. Also, one sample is misannotated (right blue dot). (D) The scatter plot for genomic sex identification generated using a contaminated dataset. The two rightmost samples are highly contaminated and are clear anomalies. The same misannotated sample is present (blue dot in the female cluster).**

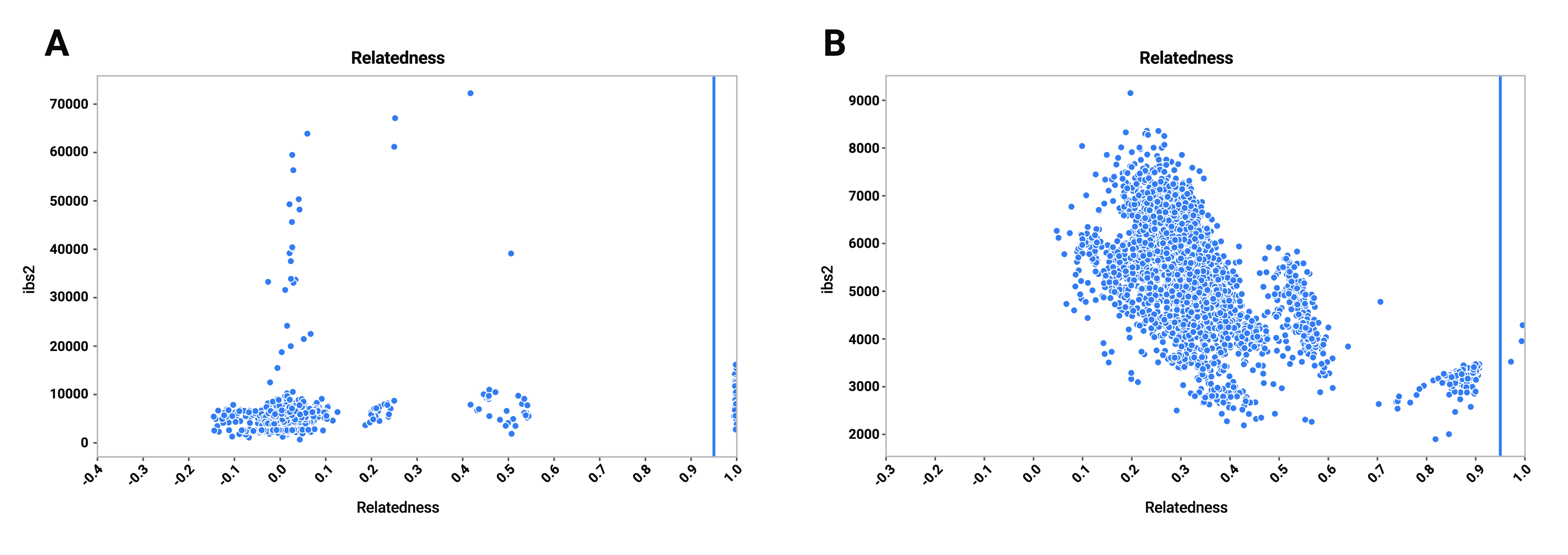

**Figure S2: Relatedness plot of a dataset (A). Each point represents a pair of samples. The cluster near 1 represents the technical replicates belonging to the same individuals. The clusters around 0.5 represent a parent-child and a sibling relationship while the cluster around 0.25 represents an aunt-nephew relationship. The clusters centered in 0 represent unrelated pairs. The points with high ibs2 are WGS pairs while the clusters with low ibs2 are WTS or WGS-WTS pairs. (B) is the same plot as (A) but generated using a highly contaminated dataset without any relationship between the samples. The unrelated cluster is centered in 0.40 because the contamination reduces the high-quality variant available for the computation of the kinship value. The highest contaminated samples are also wrongly identified as replicates/twins.**

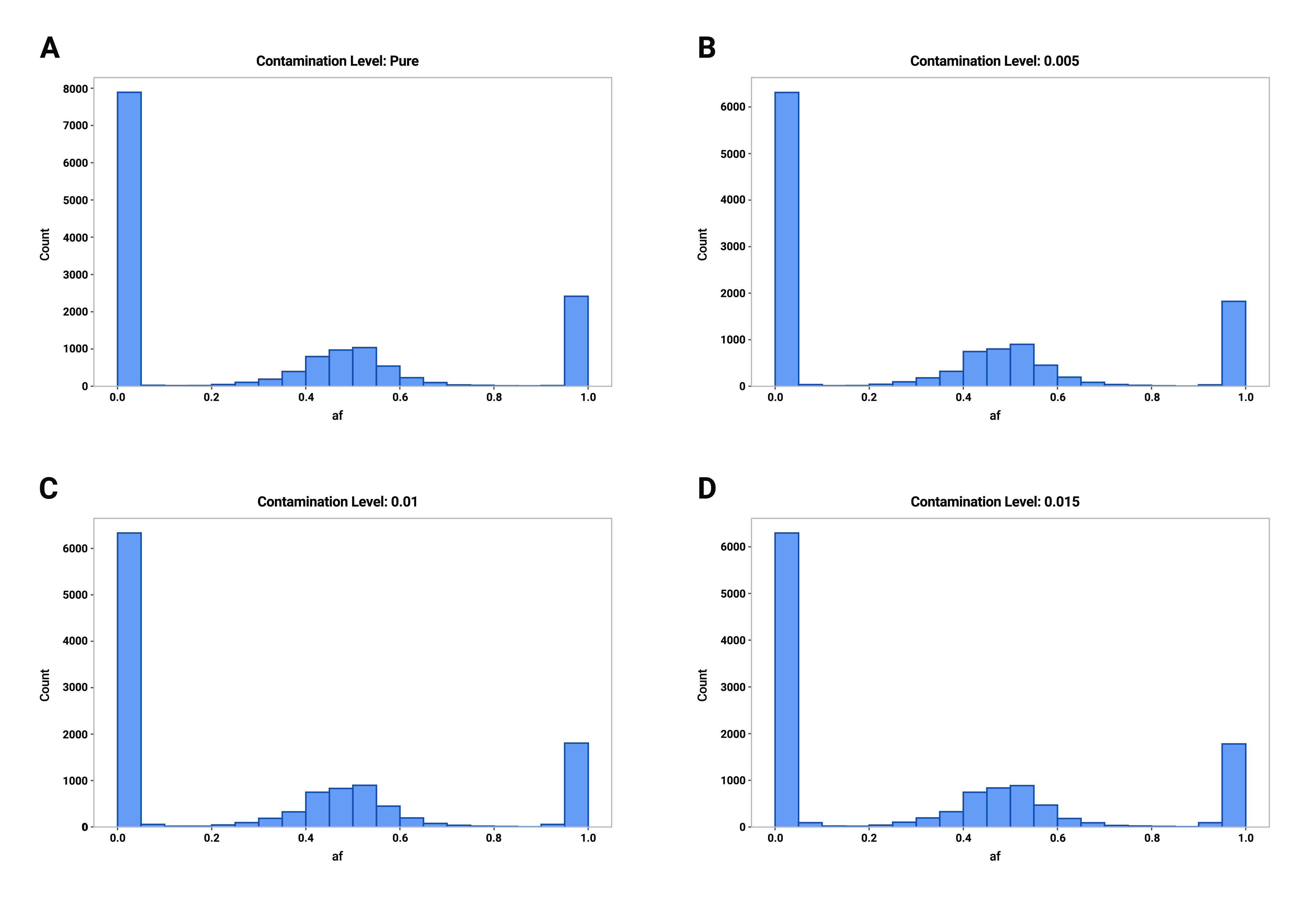

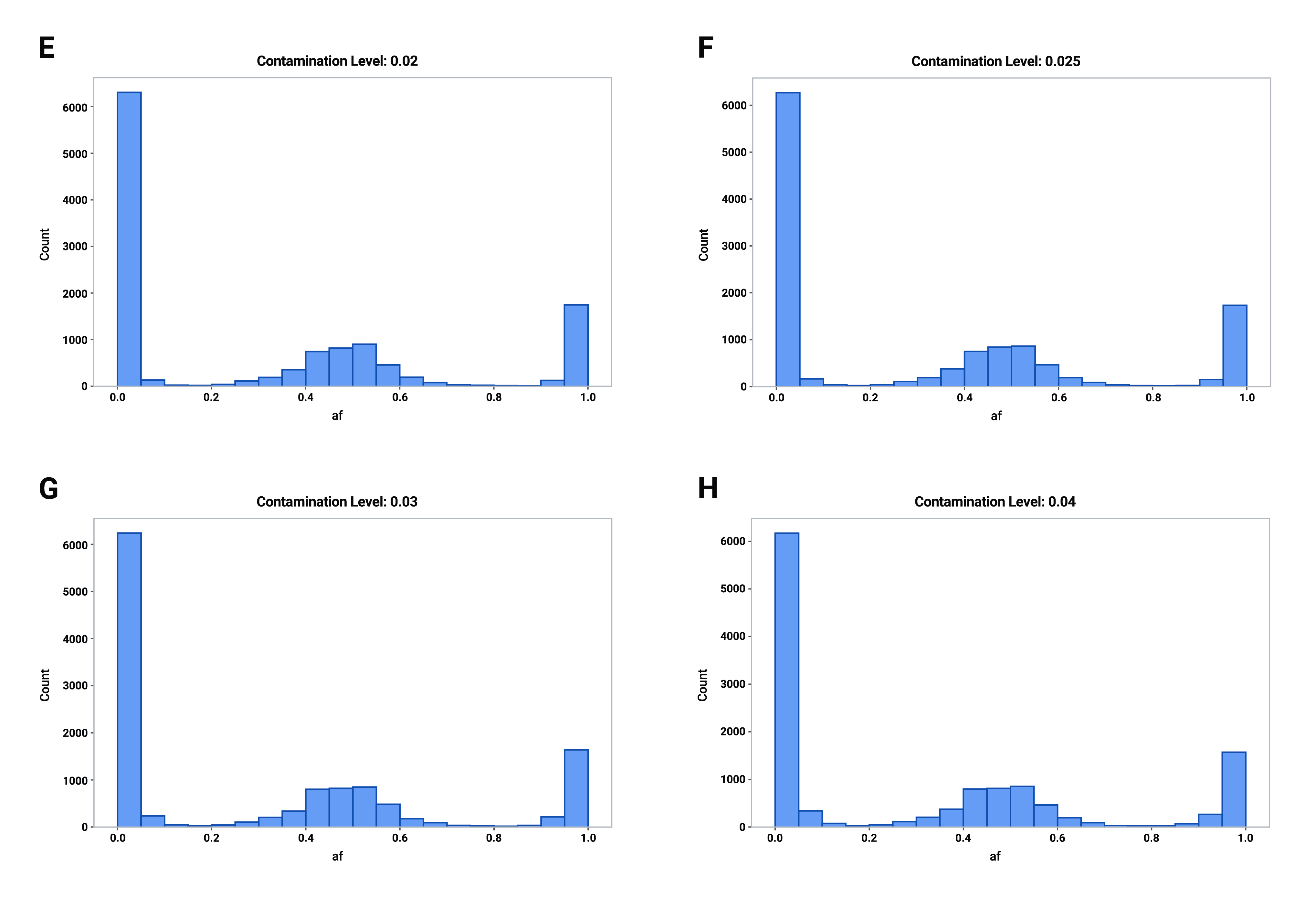

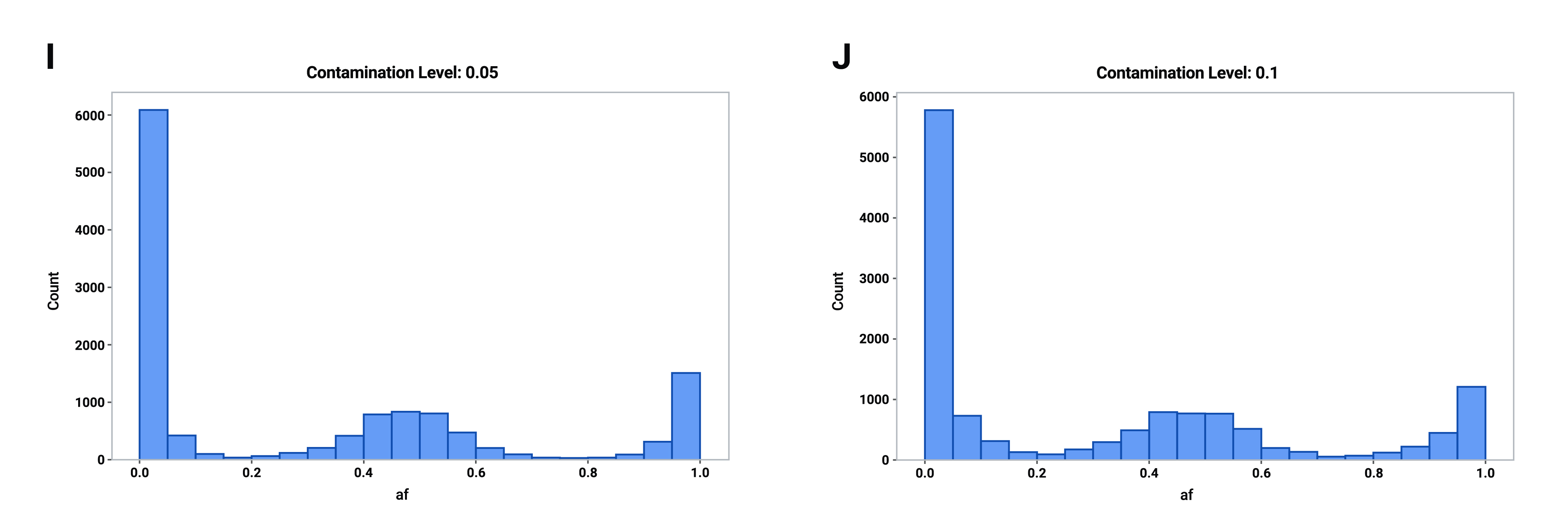

**Figure S3: Allelic fraction of the variants with an increasing level of contamination between two WTS samples. (A) pure, (B) 0.005 admixture, (C) 0.01 admixture, (D) 0.015 admixture, (E) 0.02 admixture, (F) 0.025 admixture, (G) 0.03 admixture, (H) 0.04 admixture, (I) 0.05 admixture, (J) 0.1 admixture. Overall, it’s possible to see how the number of variants in the [0.05-0.15] and [0.85-0.95] ranges increase as the contamination increases.**

**Table S1. Information on the 1000GP samples used to build the model.**

| DNA Samples | RNA Samples |
| --- | --- |
| HG00109 | HG00109 |
| HG00127 | HG00127 |
| HG00130 | HG00130 |
| HG00133 | HG00133 |
| HG00139 | HG00139 |
| HG00146 | HG00146 |
| HG00154 | HG00154 |
| HG00233 | HG00233 |
| HG00235 | HG00235 |
| HG00243 | HG00243 |
| HG00251 | HG00251 |
| HG00253 | HG00253 |
| HG00257 | HG00257 |
| HG00259 | HG00259 |
| HG00262 | HG00262 |
| HG01334 | HG01334 |
| NA06985 | NA06985 |
| NA06994 | NA06994 |
| NA07048 | NA07048 |
| NA11843 | NA11843 |
| NA11881 | NA11881 |
| NA11930 | NA11930 |
| NA11995 | NA11995 |
| NA12058 | NA12058 |
| NA12272 | NA12272 |
| NA12273 | NA12273 |
| NA12282 | NA12282 |
| NA12342 | NA12342 |
| NA12347 | NA12347 |
| NA12399 | NA12399 |
| NA12750 | NA12750 |
| NA12763 | NA12763 |
| NA12775 | NA12775 |
| NA12778 | NA12778 |
| NA12814 | NA12814 |
| NA12873 | NA12873 |
| NA12889 | NA12889 |
| NA20504 | NA20504 |
| NA20509 | NA20509 |
| NA20512 | NA20512 |
| NA20531 | NA20531 |
| NA20589 | NA20589 |
| NA20752 | NA20752 |
| NA20765 | NA20765 |
| NA20774 | NA20774 |
| NA20787 | NA20787 |
| NA20790 | NA20790 |
| NA20804 | NA20804 |
| NA20826 | NA20826 |
| NA20810 | NA20816 |

**Table S2. Contamination prediction error (Mean Absolute Error) on the RNA test dataset**

| Contamination | NgsTroubleFinder Mean Absolute Error (Test) | VerifyBamID2 Mean Absolute Error (Test) |
| --- | --- | --- |
| 0 | 0.00477 | 0.00433 |
| 0.005 | 0.00447 | 0.00518 |
| 0.01 | 0.00294 | 0.00363 |
| 0.015 | 0.00536 | 0.00540 |
| 0.02 | 0.00272 | 0.00366 |
| 0.025 | 0.00910 | 0.00856 |
| 0.03 | 0.00785 | 0.00563 |
| 0.04 | 0.00854 | 0.00816 |
| 0.05 | 0.01037 | 0.01178 |
| 0.1 | 0.02508 | 0.02940 |

**Table S3. Contamination prediction error (Mean Absolute Error) on the DNA test dataset**

| Contamination | NgsTroubleFinder Mean Absolute Error (Test) | VerifyBamID2 Mean Absolute Error (Test) |
| --- | --- | --- |
| 0 | 0.00424 | 0.00049 |
| 0.005 | 0.00195 | 0.00094 |
| 0.01 | 0.00281 | 0.00362 |
| 0.015 | 0.00353 | 0.00478 |
| 0.02 | 0.00265 | 0.00372 |
| 0.025 | 0.00508 | 0.00650 |
| 0.03 | 0.00630 | 0.00989 |
| 0.04 | 0.00694 | 0.01101 |
| 0.05 | 0.00992 | 0.01067 |
| 0.1 | 0.02272 | 0.01846 |
